# Biomechanical Modeling of a Bone Tunnel Enlargement Post ACL Reconstruction

**DOI:** 10.1101/2020.09.03.281915

**Authors:** Amirhossein Borjali, Mahdi Mohseni, Mahmoud Chizari

## Abstract

**Background:** Bone tunnel enlargement is considered as a potential problem following ACL reconstruction and can cause a fixation failure and complicate its revision surgery. This study evaluates post tibial tunnel expansion in ACL reconstruction using an interference screw.

**Methods:** A series of in-vitro experimental tests on animal bone and tissues were used to simulate post ACL reconstruction. The study believes an unbalanced lateral force can cause a local enlargement on the contact zone inside the tunnel. Grayscale X-ray images were used to assess the screw alignment inside the tunnel.

**Results:** They showed a slight misalignment between the screw and the tunnel axis as the tendon strands moved along the side of the tunnel, and the screw had partial contact with the tendon and bone along the tunnel. According to the results, increased stress in the tunnel wall causes tunnel enlargement. Although the tunnel created away from the tibial central axis produced a higher strength, it results in higher stress on the wall of the tunnel which can increase the risk of tunnel expansion.

**Conclusions:** The current study believes the use of an unguided interference screw insertion potentially increases risks of the misaligned fixation and cause a tunnel enlargement. This risk may be controlled by restricting the post-operative rehabilitation.

## 1. Introduction

Consistent graft fixation in an ACL reconstruction is a crucial element influencing the long-term success of the operation, particularly once hamstring tendon graft is applied [1]. Most of the failure in the ACL reconstruction method is due to the usage of an interfering screw to restore the graft attachment to the bone. The screw presses the graft against the bone tunnel allowing interaction occurs at the conjunction between the graft and bone. Alongside, the inserted screw keeps hold of a ligament portion inside the tunnel letting a pretension occurs on the rest of the ligament to restore the knee functionality. Conventional malfunction in the current process consists of the graft pretension loosening, cutting of the graft’s limbs by the screw threads, and withdrawal of the screw fixation due to poor bone strength or bone remodeling.

Alongside with solid fixation concerns, establishing a correct bone tunnels position at the time of surgery is a crucial phase concerning the surgical reconstruction of an ACL. Ideally, the tunnel axis should be in line with possible pullout forces to minimize lateral stresses on the surrounding bone wall. It is expected an unbalanced lateral force causes local damage or deformation on the contact zone on the boney wall inside the tunnel. As a result, the unbalanced deformation on the tunnel wall, in the post-operation period, can cause an enlargement inside the bony tunnel which can end up with a failure in fixation. Postoperative bone tunnel expansion has preceded to a revised ACL surgery [2], perhaps because of bone stress deprivation around the tunnels [3]. Although a variety of tunnel locations has been formed, the locations have not been assessed from a mechanical viewpoint to diminish negative effects.

Many surgeons routinely use bio- or nonbio-absorbable aperture interference screw fixation for ACL reconstruction. However, in all types of interference screws, the postoperative tunnel enlargement problem exists [4]. Depends on the level of local enlargement, it has been shown that the tunnel enlargement can affect the outcome of ACL reconstruction and complicate its revision surgery [5].

ACL fixation at the tibial side is more problematic than at the femoral side as the armed forces applied to the reconstructed tissue is parallel to the tibial tunnel axis [6]. On the other hand, the tibial bone quality from the mechanical point of view is weaker compare to the femoral bone [1]. If a quadruple tendon graft strand is used as a graft, the four-tailed end of the tendon graft is trickier to be secured at the tibial tunnel. By the aid of whip stitching, the looped portion of the connective tissue can be easily bundled into a compressed shape and pulled through the tibial tunnel and hanged in the femoral tunnel using an Endo-button. While the four loose ends the graft should be secured using an interference screw or other fixation devices.

In the literature, there are studies about bone tunnel enlaegement after ACL reconstruction evaluating the level of tunnel expansion post operation using a variety of interference screws [7–9]. However, it is well-known that the tunnel local or cross-sectional region could widen in the first 12 weeks post-operation [4]. Furthermore, the researches on both tibial and femoral tunnel growth shown that after auto-graft ACL reconstruction the mechanical instabilities are the main contributed factor [5, 8].

The current study aims to investigate the tunnel expansion following an ACL reconstruction when an interference screw is used and as a fixation implant. The research program includes a precise examination of an interference screw fixation through an experimental study. Nevertheless, this work uses porcine tibial bone and bovine flexor tendon for its experimental evaluation, the animal bone has been used in a previous study on graft fixation and has been shown to give results that are not substantially different from those which were found with young human bones [1].

## 2. Materials and methods

Fresh porcine tibia bone and bovine tendon were used in the experimental part of this study. The harvesting procedure and the biomechanical test protocol were approved by the local clinical officer. The harvested tibia bone and tendon were trimmed from supporter muscle fibers and surrounding connective tissues, packed, and kept frozen at −20°C in sealed bags for no more than two months. The preservation policy is proposed for samples intended for the in vitro testing procedure of the ligament reconstructions. They have proven that they do not affect the bones’ mechanical properties [10].

On testing day, the bone and tendons were exposed to room temperature (tendon for 2-4 hours, porcine tibia bone for 6-8 hours). All the specimens held moist with water during the samples’ preparation, fixation methods, and mechanical testing. The tendon graft diameter was chosen to permit a close fit passage into the bone tunnel (about 0.5 mm less than the tunnel diameter). The tendon strands were trimmed to the identical size. According to the technique described in a study by Chizari et al. [1, 11], two same sized tendon strands with a total length of 160 mm were coupled (double-stranded) and folded in half to build a four-strand bundle with a total 80 mm graft length.

While preserving a steady tension on all four strands, the graft was sutured at the free end for 25 mm with No. 2 vicryl suture using a whip stitch. The size of the bundle was measured with a graft sizing scale to make sure that the diameter was the same for all sections of the specimen. Considering the procedure described by Chizari et al. [1, 12], the tibial bone was prepared for tunneling using a universal drill pack. The bone tunnel was placed in a normal surgical orientation in the tibia bone. Firstly, a small hole to take a guidewire was drilled into the tibia bone in the desired location with the aid of an ACL guide instrument and then the main tunnel was drilled using a 10 mm conventional cannulated drill bit (Fig. 1. a). The sharp corner of the tunnel was then smoothed using a special dilator after tunneling. After the tunnel was drilled the looped end of the bundle was dragged throughout the tunnel with the help of the surgical guide, leaving the tightly sutured four-tailed end of the bundle inside the bone tunnel (Fig. 1. b).

**Fig. 1.**
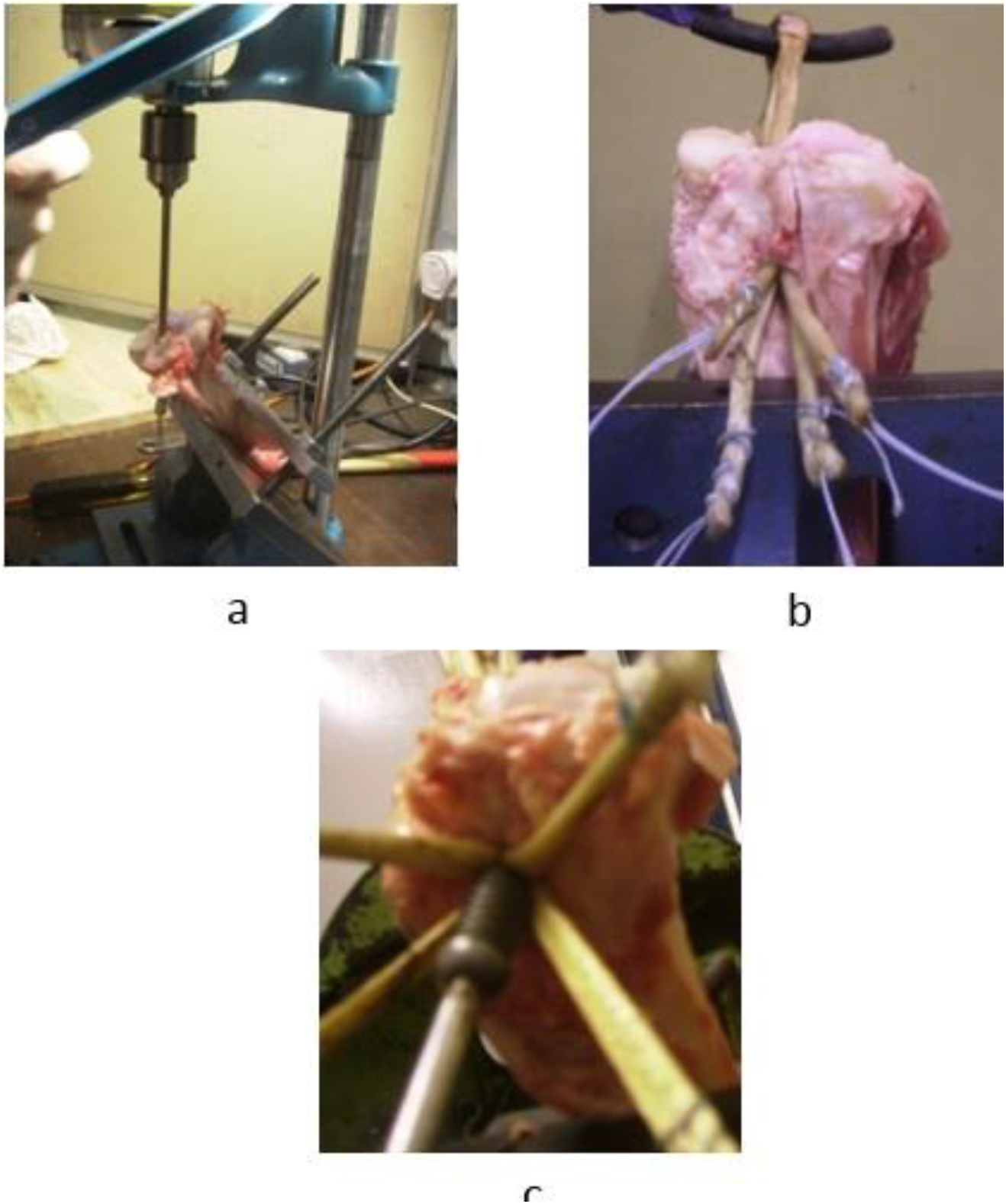
a. Practical stages of creating a bone channel, b. Insertion the graft, c. Inserting the interference screw into a porcine tibia bone tunnel

The tendon bundle was located through the tibia bone tunnel with 30 to 35 mm of the unsutured, looped part of the tendon sticking out from the proximal opening of the bone tunnel. This length related to the length of the normal ACL [13]. By maintaining the four-tailed end of the tendon bundle, a 10 mm diameter interference screw was screwed into the entrance (anterior aspect) of the tunnel between the tendon strands and moved forward until its tip reached the proximal bone-tunnel opening (Fig.1 c).

A general biomechanical test, i.e., tensile test as described previously [14–16], was performed on the bone and tendon to observe the placement of the graft and screw in the bone tunnel after the mechanical loading. The loop portion of the tendon hanged with the crosshead of the testing machine, and the bone was secured using custom-made grips in the testing machine (Fig. 2). The sample’s structure was preconditioned by a cycling load at 1 Hz from 10 to 50 N for ten cycles, followed by a cyclic loading from 50 to 250 N at 1 Hz for 500 cycles. After that, if the structure was not failed in previous stages, a load-to-failure test was carried out at a rate of 20 mm/min [14, 16]. Failure was characterized either by tendon rupture or when significant slip-up occurs at fixation [17, 18].

**Fig. 2.**
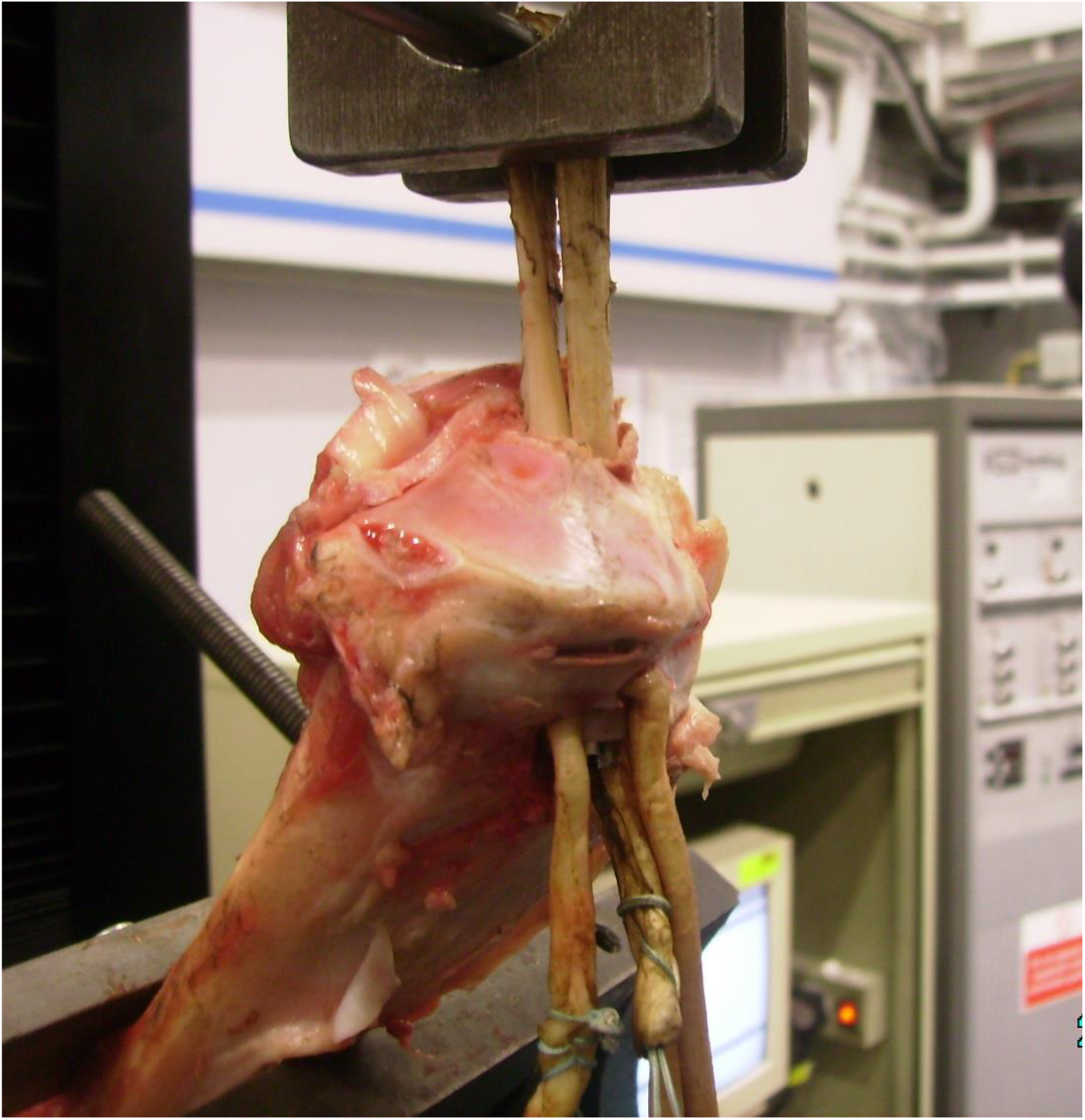
Pull out test on a porcine tibia bone specimen

## 3. Results

Several bovine and porcine tibia bone specimens were prepared using the procedure described earlier. To examine the placement of the interference screw inside the tunnel, some of the specimens were examined using X-ray scanning. Furthermore, to evaluate the contact zone between the screw and the tendon limbs or bone tunnel, some of the specimens were frozen and then cut in slices along the axial direction, and cross-sections were closely monitored (Fig. 3. a and b).

**Fig. 3.**
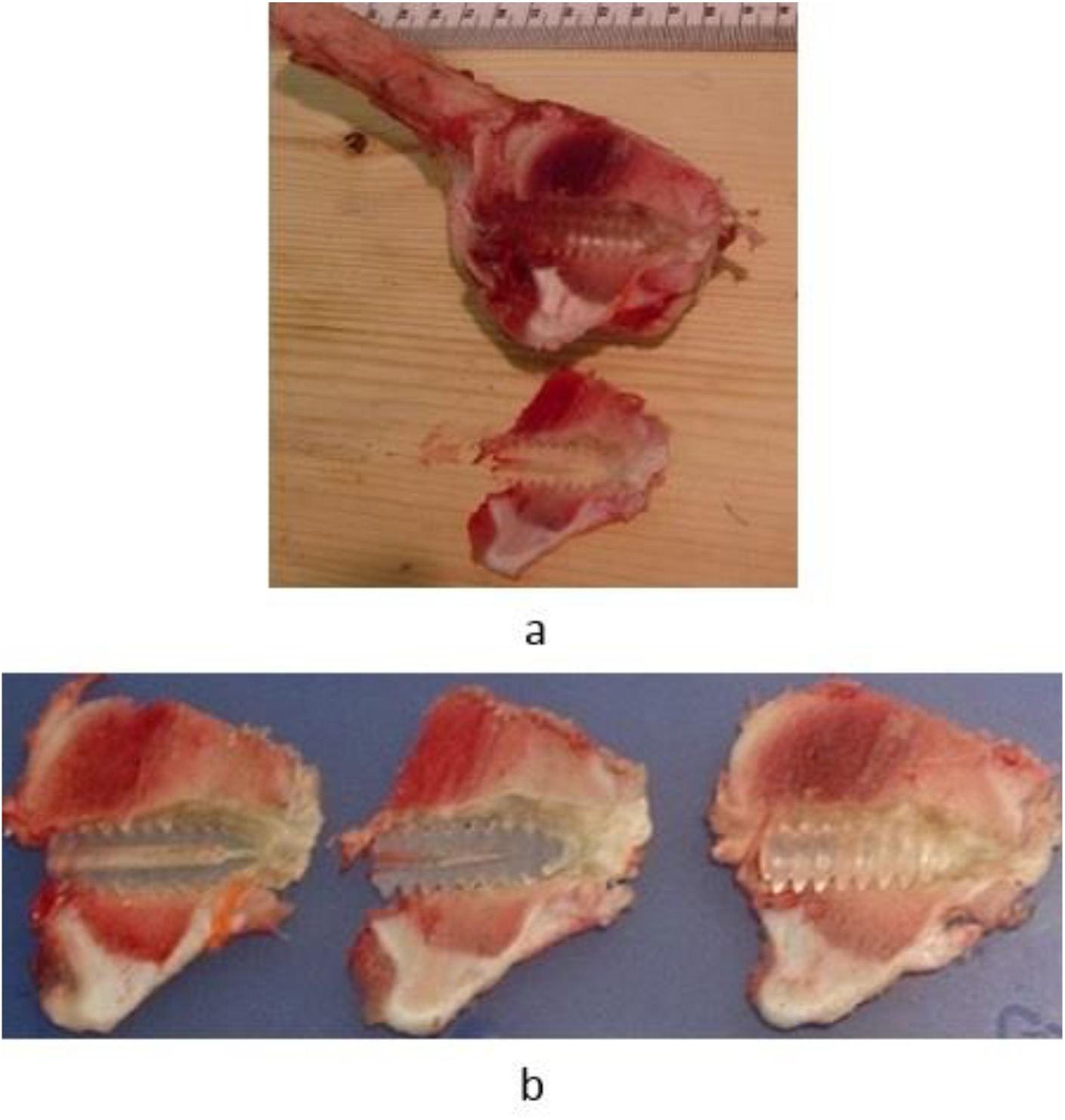
a. Sectioning of a porcine tibia bone along the axial direction of the tunnel and b. Observing the detail of the screw placement into the tunnel. A bio-absorbable screw was used to fix the tendon inside the bone tunnel.

The screw placement inside the tunnel is essential, as any postoperative change in the screw position will affect the mechanical stability of the fixation. A situation was expected that the screw would symmetrically place among the bundle’s strands allied with the axis of the tunnel, but in rehearsal when the screw inserts, it twists and forces the tendon strands aside from the axis of the tunnel and the tendons move at incidental route so that the strands form an unsymmetrically arrangement. This was confirmed by sectioning the frozen specimen along the axial direction and agrees with findings by other researchers [19] (Fig. 3. a and b).

Further examination of slices showed that the screw is partially in contact with tendon limbs and the boney wall (Fig. 3. b). Close examination showed the screw threads were cut into the bone causing local damage on the bone (Fig. 4. a and b). Examination of the slices showed that the screw also become misaligned with the axis of the tunnel. This examination showed that the sideways motion of the screw within the tunnel triggers an eccentric placement and excites the enlargement (Fig. 4. c).

**Fig. 4.**
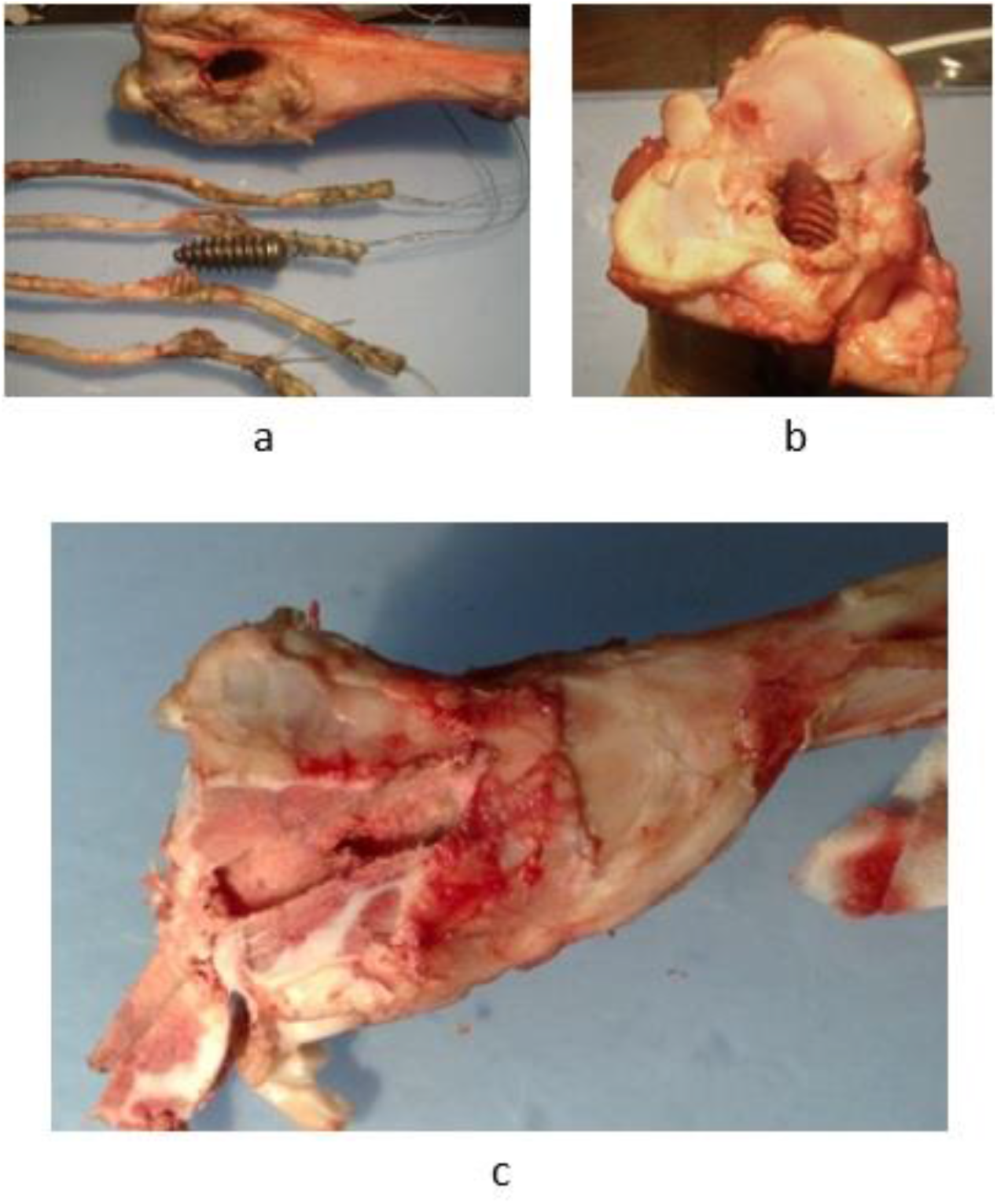
a. A typical specimen with failure at its fixation, b. The effect of the screw cut on the bone, c. The lateral movement of the screw inside the tunnel causes an eccentric position and enlargement

Generally, tunnels created away from the tibial central axis produced lower axial stress on the bone tunnel and resulted in higher pullout strength. Of course, this is anticipated as less rigid bone exists in the central region, and the pullout loads apply un-axially. However, a danger may exist at the tunnel where the wall is too close to the bone corner. Although this may increase the fixation strength as the tunnel created in the region close to the cortical bone, it may increase the risk of fracture during the screw insertion as the tunnel surrounding wall is thin in this region. Furthermore, un-axial or unbalanced pullout loads create lateral stress and mechanical instability on the tunnel wall which can lead to enlargement. As a result, tunnel enlargement will resonate with the stress in the lateral tunnel wall and cause more enlargement and damage to the bone tunnel.

To validate the finding from slices, an X-ray scanner was used to investigate the outplacement of the screw inside the tunnel (Fig. 5. a and b). For each sample, an X-ray scan was obtained on sagittal and coronal planes. To get a clear image on the true site of the bone tunnel, the scan was focusing on the proximal tibia bone approximately 100 mm distal from the articular surface.

**Fig. 5.**
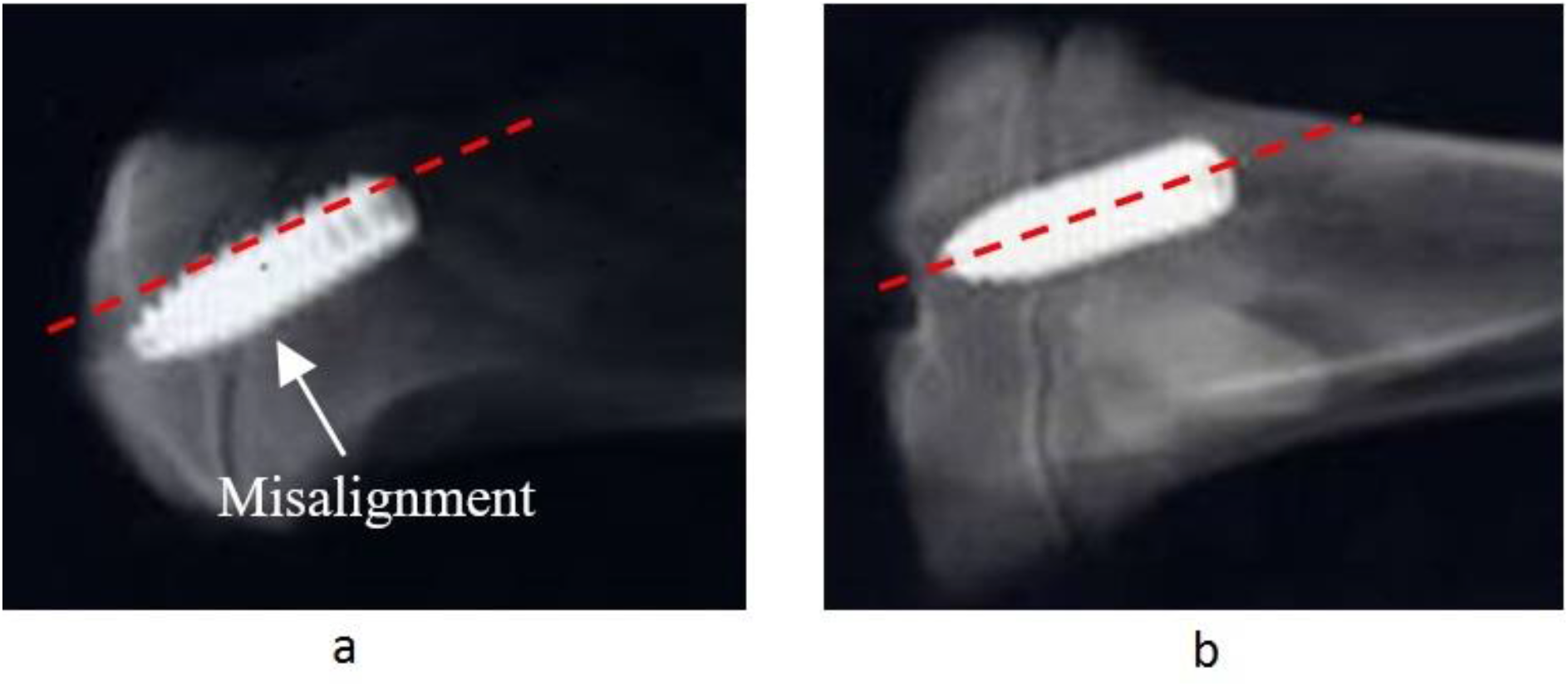
a. Sagittal and b. coronal views of X-ray scans of a porcine tibia bone screw fixation.

Following the X-ray examination, the alignment of the fixation with the bone tunnel was assessed. The X-ray images proved the existence of a misalignment between the screw and the tunnel axis, at least in one of the planes. This means during the screw insertion process the tendon strands pushed aside partially and the screw was tilted to touch the bone tunnel. Therefore, the screw had only partial contact with the tendon and a partial contact with the bone along the tunnel.

## 4. Discussion

Though several investigators have demonstrated that bone minor tunnel expansion does not seem to involve graft laxity or failure rates adversely, in case of significant enlargement and failure, a review or revision surgery could be cruelly hampered by creating of a larger tunnel [5]. The tunnel expansion, which exists by the joint movement, frequently maximizes the middle district of the tunnel [7, 20]. The event etiology still is not certain, but it is unquestionably a multifactorial issue, consist of different parameter e.g. mechanical and biological factors. It expected many parameters to be involved such as mechanical elements containing tunnel positioning errors and graft-tunnel relative micro-motion [4]. Early mobilization is the next possible parameter, which aggravates the elements’ micro-motion before the graft been properly healed [21].

An interfering screw insertion pushes the graft against the tunnel due to its high compressive stiffness of the implant contrasted to the cancellous bone [22]. Furthermore, eccentric positioning of the interfering screw into the tunnel may initiate a tunnel enlargement and possible cyclical loads may excite enlargement and lead it to failure.

In the current experimental study, an identical surgical method was used to construct the fixation using tendon graft and interference screw. The samples then tested using a cyclical pullout test. The average calculated enlargement for the tested samples was 35% and can be considered as a minor enlargement as the suggested threshold for tunnel enlargement is 50% [20].

It is interesting to note that, no tunnel enlargement can be observed at the distal end of the tibial tunnel, regardless of tunnel morphology. This is likely due to the tunnel is free of any tendon material in this region. It is noteworthy that the tunnel enlargement rates from 2-dimensional plain radiographs. Commonly, calculated enlargement is determined either by measuring the tunnel cross-section or, more frequently, it is demonstrated as a percentage escalation of the tunnel diameter. As an example, a 1 mm escalation in a previously drilled 10 mm tunnel is acted for a 10% increase in diameter while it is indicating 17.4% enlargement on the tunnel cross-section. It is therefore extra care expected in interpreting reports for tunnel expansion when simple radiograph evaluations have been used [23].

The motion of the tendon bundle inside the tunnel shown to cause a notable tunnel enlargement [24]. Two ways of motion have been explained: axial motion, which relates to its bungee or cyclical effect, and radial motion, which is at the right angle to the axis of the tunnel and causes windshield wiper effect [9]. Although both types of motions may affect the healing of the fixation, the radial motion of the graft can cause a gap between bone and graft. If the gap becomes too large, non-union or delayed union may happen even with enough fixation. The gap may be reduced at the tunnel entrance by redirecting the force applies to the graft [25]. This means the tunnel angle must be carefully chosen before reconstruction starts. The graft tissue fixation close to the articular entrance of the tunnels does influence graft-bone tunnel pressure; an interference screw placed on the side of the tunnel that encounters the orientation of the force vector can apply a converse force on the graft tissue. In this way, the graft migration in the direction of the force will be enhanced [9]. Redirecting forces may be an essential factor that causes bone tunnel enlargement as the reconstruction is stressed mechanically. The extent of redirecting loads depends on the angle of the tunnel, graft tensile load, and the loads apply from the fixation device inside the tunnel. Since the graft tension at different positions of the knee varies, redirecting forces may be higher when the tunnel is steeper at the extended knee. The femoral bony tunnel, particularly in the sagittal plane, is an example of this problem. A potential resolution to this issue can be to change the tunnel angulation. Rigid fixation is another crucial that can restrain redirecting loads.

It has been shown that argumentative rehabilitation has corresponded to the bone tunnel enlargement [7]. Immobilization would undoubtedly diminish the forces, which can cause a enlargement, but it will slow down the rehabilitation process. Hence, instead of immobilization, controlled exercising in the safe range of motion [7] can be recommended. It is suggested to use organic implant-less ACL reconstruction methods such as BASHTI technique to prevent the tunnel enlargement phenomenon [26].

## Conclusion

The current study examines the bone and tendon graft interaction in a tunnel when an ACL reconstructed performs at the knee. The study emphasis was on ACL reconstruction when interference screws combine with hamstring graft is used. A close look at the fixation site for possible bone tunnel expansion under a full load-bearing post-operation was the concern of the study. A series of in-vitro experimental studies were performed using animal stuff and the interaction of soft tissue ant implant at the tunnel was observed. The experimental observation revealed that by inserting the interference screw into the tunnel, the screw twist and pushes the tendon away and may penetrate the tunnel wall. It was also indicated the side movement of the implant within the tunnel might be positioned eccentrically and excite the bone tunnel enlargement. Redirecting force may also cause bone tunnel enlargement. The magnitude of the force mainly depends on the tunnel angle and can be controlled by changing the angle. The study believes a misaligned interference screw potentially increases the risk of a tunnel enlargement. However, the risk of widening of the tunnel can be reduced by controlling the screw insertion at the time of operation and a restricted rehabilitation post-operation.

## Reference

1. Chizari M, Wang B, Barrett M, Snow M (2010) BIOMECHANICAL TESTING PROCEDURES IN TENDON GRAFT RECONSTRUCTIVE ACL SURGERY. Biomed Eng Appl Basis Commun 22:427–436. https://doi.org/10.4015/S1016237210002195

2. Rizer M, Foremny GB, Rush A, et al (2017) Anterior cruciate ligament reconstruction tunnel size: causes of tunnel enlargement and implications for single versus two-stage revision reconstruction. Skeletal Radiol 46:161–169. https://doi.org/10.1007/s00256-016-2535-z

3. Shimizu R, Adachi N, Ishifuro M, et al (2017) Bone tunnel change develops within two weeks of double-bundle anterior cruciate ligament reconstruction using hamstring autograft: A comparison of different postoperative immobilization periods using computed tomography. Knee 24:1055–1066. https://doi.org/10.1016/j.knee.2017.06.013

4. Marchant MHJ, Willimon SC, Vinson E, et al (2010) Comparison of plain radiography, computed tomography, and magnetic resonance imaging in the evaluation of bone tunnel widening after anterior cruciate ligament reconstruction. Knee Surg Sports Traumatol Arthrosc 18:1059–1064. https://doi.org/10.1007/s00167-009-0952-4

5. Lopes OVJ, de Freitas Spinelli L, Leite LHC, et al (2017) Femoral tunnel enlargement after anterior cruciate ligament reconstruction using RigidFix compared with extracortical fixation. Knee Surg Sports Traumatol Arthrosc 25:1591–1597. https://doi.org/10.1007/s00167-015-3888-x

6. Petre BM, Smith SD, Jansson KS, et al (2013) Femoral cortical suspension devices for soft tissue anterior cruciate ligament reconstruction: a comparative biomechanical study. Am J Sports Med 41:416–422. https://doi.org/10.1177/0363546512469875

7. Leonardi AB de A, Duarte Junior A, Severino NR (2014) Bone tunnel enlargement on anterior cruciate ligament reconstruction. Acta Ortop Bras 22:240–244. https://doi.org/10.1590/1413-78522014220500338

8. Ge Y, Li H, Tao H, et al (2015) Comparison of tendon-bone healing between autografts and allografts after anterior cruciate ligament reconstruction using magnetic resonance imaging. Knee Surg Sports Traumatol Arthrosc 23:954–960. https://doi.org/10.1007/s00167-013-2755-x

9. Lee DW, Lee JW, Kim SB, et al (2017) Comparison of Poly-L-Lactic Acid and Poly-L-Lactic Acid/Hydroxyapatite Bioabsorbable Screws for Tibial Fixation in ACL Reconstruction: Clinical and Magnetic Resonance Imaging Results. Clin Orthop Surg 9:270–279. https://doi.org/10.4055/cios.2017.9.3.270

10. Kondo E, Merican AM, Yasuda K, Amis AA (2010) Biomechanical comparisons of knee stability after anterior cruciate ligament reconstruction between 2 clinically available transtibial procedures: anatomic double bundle versus single bundle. Am J Sports Med 38:1349–1358. https://doi.org/10.1177/0363546510361234

11. Chizari M, Alrashidi M, Alrashdan K, et al (2013) Mechanical Aspects of an Interference Screw Placement in ACL Reconstruction. In: IWBBIO. Granada

12. Chizari M, Snow M, Wang B (2011) Post-operative assessment of an implant fixation in anterior cruciate ligament reconstructive surgery. J Med Syst 35:941–947. https://doi.org/10.1007/s10916-010-9514-z

13. Taylor KA, Cutcliffe HC, Queen RM, et al (2013) In vivo measurement of ACL length and relative strain during walking. J Biomech 46:478–483. https://doi.org/10.1016/j.jbiomech.2012.10.031

14. Snow M, Cheung W, Mahmud J, et al (2012) Mechanical assessment of two different methods of tripling hamstring tendons when using suspensory fixation. Knee Surgery, Sport Traumatol Arthrosc 20:262–267. https://doi.org/10.1007/s00167-011-1619-5

15. Chizari M, Wang B, Snow M, Barrett M (2010) Mechanical aspects of a single-bundle tibial interference screw fixation in a tendon graft ACL reconstruction. Biomed Eng Appl Basis Commun 22:. https://doi.org/10.4015/S101623721000202X

16. Chizari M, Snow M, Cheung W, et al (2012) Relative motion of tendon limbs in a loop tendon graft. Biomed Eng Appl Basis Commun 24:1–5. https://doi.org/10.1142/S1016237212500408

17. Yang D-L, Cheon S-H, Oh C-W, Kyung H-S (2014) A comparison of the fixation strengths provided by different intraosseous tendon lengths during anterior cruciate ligament reconstruction: a biomechanical study in a porcine tibial model. Clin Orthop Surg 6:173–179. https://doi.org/10.4055/cios.2014.6.2.173

18. Higano M, Tachibana Y, Sakaguchi K, et al (2013) Effects of tunnel dilation and interference screw position on the biomechanical properties of tendon graft fixation for anterior cruciate ligament reconstruction. Arthrosc J Arthrosc Relat Surg Off Publ Arthrosc Assoc North Am Int Arthrosc Assoc 29:1804–1810. https://doi.org/10.1016/j.arthro.2013.07.263

19. Saithna A, Chizari M, Morris G, et al (2015) An analysis of the biomechanics of interference screw fixation and sheathed devices for biceps tenodesis. Clin Biomech 30:551–557. https://doi.org/10.1016/j.clinbiomech.2015.04.006

20. Streich NA, Reichenbacher S, Barié A, et al (2013) Long-term outcome of anterior cruciate ligament reconstruction with an autologous four-strand semitendinosus tendon autograft. Int Orthop 37:279–284. https://doi.org/10.1007/s00264-012-1757-5

21. Panariello RA, Stump TJ, Allen AA (2017) Rehabilitation and Return to Play Following Anterior Cruciate Ligament Reconstruction. Oper Tech Sports Med 25:181–193. https://doi.org/10.1053/j.otsm.2017.07.006

22. Xu Y, Ao Y, Wang J, et al (2011) Relation of tunnel enlargement and tunnel placement after single-bundle anterior cruciate ligament reconstruction. Arthrosc J Arthrosc Relat Surg Off Publ Arthrosc Assoc North Am Int Arthrosc Assoc 27:923–932. https://doi.org/10.1016/j.arthro.2011.02.020

23. Athapattu M, Amir H, Saveh AH, et al (2013) Accuracy of Measuring Methods on the Femoral Head. Procedia Eng 68:. https://doi.org/10.1016/j.proeng.2013.12.151

24. Silva A, Sampaio R, Pinto E (2010) Femoral tunnel enlargement after anatomic ACL reconstruction: a biological problem? Knee Surg Sports Traumatol Arthrosc 18:1189–1194. https://doi.org/10.1007/s00167-010-1046-z

25. Xu H, Zheng R, Ying J (2017) Bone Tunnel Impaction Reduced the Tibial Tunnel Enlargement. Open Med (Warsaw, Poland) 12:99–106. https://doi.org/10.1515/med-2017-0016

26. Moeinnia H, Nourani A, Borjali A, et al (2020) Effect of Geometry on the Fixation Strength of Anterior Cruciate Ligament Reconstruction Using BASHTI Technique. J Knee Surg. https://doi.org/10.1055/s-0040-1716371

